# Genome-wide signatures of genetic variation within and between populations – a comparative perspective

**DOI:** 10.1101/104604

**Authors:** Nagarjun Vijay, Matthias Weissensteiner, Reto Burri, Takeshi Kawakami, Hans Ellegren, Jochen B. W. Wolf

## Abstract

Genome-wide screens of genetic variation can reveal signatures of population-specific selection implicated in adaptation and speciation. Yet, unrelated processes such as linked selection arising as a consequence of genome architecture can generate comparable signatures across taxa. To investigate prevalence and phylogenetic stability of linked selection, we took a comparative approach utilizing population-level data from 444 re-sequenced genomes of three avian clades spanning 50 million years of evolution. Levels of nucleotide diversity (π),population-scaled recombination rate (ρ), genetic differentiation (F_ST_, PBS) and sequence divergence (D_xy_) were remarkably similar in syntenic genomic regions across clades. Elevated local genetic differentiation was associated with inferred centromere and sub-telomeric regions. Our results support a role of linked selection shaping genome-wide heterogeneity in genetic diversity within and between clades. The long-term conservation of diversity landscapes and stable association with genomic features make the outcome of this evolutionary process in part predictable.

## INTRODUCTION

Understanding the processes governing heterogeneity of genome-wide diversity has been a long-standing goal in evolutionary genetics (Ellegren & Galtier 2016) and is of central importance to adaptation and speciation research (Seehausen et al. 2014) A plethora of recent studies quantifying genetic variation within and between natural populations share the central observation of strong heterogeneity in genome-wide distribution of genetic diversity (Seehausen et al. 2014). Despite commonality in patterns seen across a wide range of taxa, elucidating the underlying processes remains challenging (Wolf & Ellegren 2016). Regions of reduced genetic diversity generally coinciding with elevated differentiation (Charlesworth 1998) can be interpreted in terms of reproductive isolation resulting from divergent selection against homogenizing gene flow in the context of adaptation and speciation ('speciation islands') (Nosil & Feder 2013). Yet, linked selection either in the form of genetic hitch-hiking (Smith & Haigh 1974) or background selection (Charlesworth et al. 1993) can likewise introduce heterogeneity in genomic differentiation by local reduction of the effective population size (N_e_), even in the absence of divergent selection and gene flow (Cutter & Payseur 2013; Cruickshank & Hahn 2014). Discriminating between these scenarios is complicated by the fact that a multitude of intrinsic and extrinsic factors influence genome-wide patterns of diversity and differentiation (Strasburg et al. 2012)

Several ways forward have been suggested to isolate the underlying processes. *Functional validation* of candidate genes flagged during genome scans can provide valuable, independent information (Kronforst & Papa 2015). *Theoretical models* provide useful null expectations to compare with empirical patterns (Bank et al. 2014). *Experimental evolution* studies (Dettman et al. 2007) or manipulative experiments in natural populations (Soria-Carrasco et al. 2014) allow studying the link between selection and genomic patterns of genetic diversity under controlled conditions. *Micro-level comparative population approaches* leveraging information from spatiotemporal contrasts between populations (speciation continuum' (Seehausen et al. 2014)) help disentangle the effects of linked selection unrelated to speciation from those thought to contribute to reproductive isolation (Wolf & Ellegren 2016). Within species and among closely related species, however, a substantial fraction of genetic variation is shared by ancestry impeding inference.

Here, we propose a *macro-level comparative approach* extending comparisons of genome-wide diversity beyond closely related taxa to phylogenetically distant clades, where lineage sorting has since long been completed. Co-variation in the landscape of genetic diversity across clades can only be mediated by shared processes independent of population-specific selection pressures. One candidate process is the mutation rate which is known to vary across the genome (Hodgkinson & Eyre-Walker 2011). Another is linked selection, where local reduction in effective population size (N_e_) through selection depends on the rate of local recombination likewise varying across the genome (Cutter & Payseur 2013). While support for a role of mutation rate in modulating the level of genetic variation across the genome has been low (Cutter & Payseur 2013) there is increasing evidence for linked selection as a key process (Rockman et al. 2010; Slotte 2014; Burri et al. 2015). With evidence for long-term conservation of broad-scale recombination rates (Auton et al. 2012; Kawakami et al. 2014) the comparison of summary statistics reflecting local N_e_ in syntenic regions among clades holds information on the role of linked selection for each species (**Fig. 1**).

**Figure 1.**
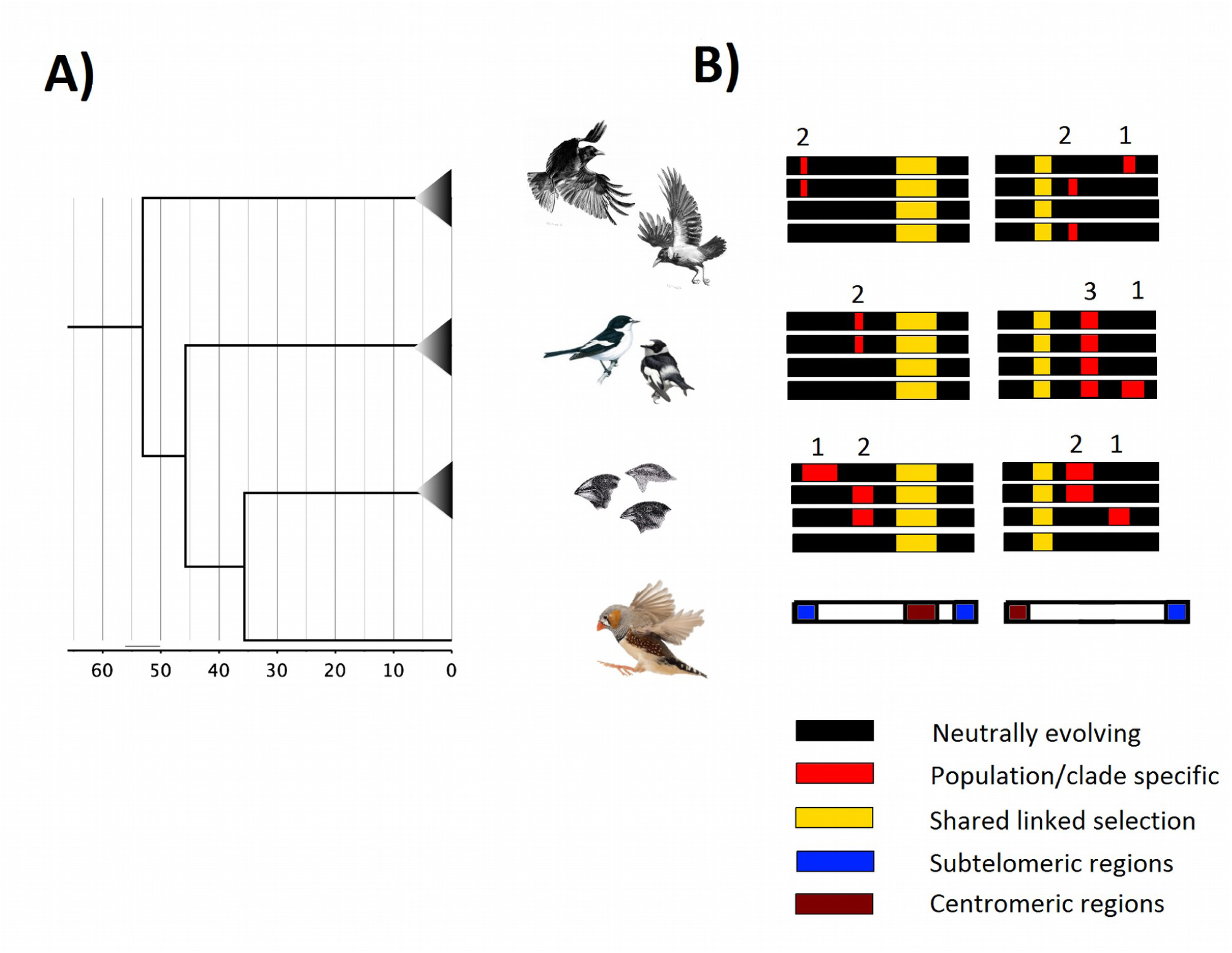
**Study design A)** Dated phylogenetic reconstruction of all clades used in this study. Note that for each focal taxon (crows, flycatchers and Darwin's finches) a large number of individuals from several populations and (sub-)species have been used including 120 Darwin's finch genomes (Lamichhaney et al. 2015), 200 genomes of *Ficedula* flycatchers (Burri et al. 2015) and 124 genomes of crow from the genus *Corvus* (Vijay et al. 2016). **B)** Rationale of the study exemplified for two schematic chromosomes and four populations of each clade. We calculated summary statistics of genetic diversity within and between populations of each clade indicative of regional effective population size (N_e_). Genomic regions with reduced local N_e_ indicating (1) putative directional selection in single populations or (2) divergent selection against gene flow in population pairs are symbolized in red. Due to shared recent common ancestry signatures reflecting past selection reducing N_e_ may occur across a several populations (3). Demographic perturbation increasing variance in N_e_ across the genome may likewise generate extreme values of N_e_ for specific populations. Regional reduction in local N_e_ across clades, where lineage sorting has been completed, is depicted in yellow. Shared signatures are best explained by linked selection affecting syntenic regions similarly in all clades. Genomic regions evolving predominantly according to neutrality are shown in black.

We utilized genome-wide re-sequencing data from several populations or (sub)-species of three distantly related avian species complexes - Darwin's finches, *Ficedula* flycatchers and *Corvus* crows- with split times beyond the expected time for complete lineage sorting (**Fig. 1, Supporting Information**). For each population and population comparison within clades, we quantified genetic summary statistics in syntenic windows of 50 kb in size. Summary statistics were chosen to be reflective of the local effective population size (N_e_) of a genomic region: population-scaled recombination rate ρ (∼N_e_r), nucleotide diversity π (∼N_e_µ), a measure of genetic differentiation F_ST_(∼1/(1+N_e_ (m+µ)) (where mutation rate µ can generally be neglected if migration rate m>>µ), the related population branch statistic (PBS) accounting for non-independence of population comparisons, and sequence divergence D_xy_ (∼N_e_µ+µt). The only parameter shared by these statistics is N_e_; hence, co-variation of all statistics in syntenic regions would indicate shared processes affecting local N_e_ alike in the investigated populations.

## MATERIAL AND METHODS

### Clades

As subject for this investigation we chose populations and (sub)-species from three phylogenetically divergent clades: Darwin's finches of the genera *Geospiza, Certhidea and Platyspiza.*, flycatchers of the genus Ficedula (*F. albicollis, F. hypoleuca, F. semitorquata* and *F. speculigera)* and crows of the genus *Corvus* including the American crow *C. brachyrhynchos* and several taxa from the *Corvus (corone) spp.* species complex (Vijay et al. 2016). Functionally annotated genome assemblies with high sequence contiguity are available for one representative each of *Ficedula* flycatchers (*F. albicollis*, genome size: 1.13, scaffold/contigN50=7.3/410kb, NCBI accession number: GCA_000247815.2) (Ellegren et al. 2012; new chromosome build Kawakami et al. 2014) and for one hooded crow specimen (*Corvus (corone) cornix*, genome size: 1.04Gb, scaffold/contig N50=16.4Mb/94kb, National Center for Biotechnology Information (NCBI) accession number: GCA_000738735.1) (Poelstra et al. 2014, 2015). The assembly of the medium ground finch *G. fortis* is of comparable size (1.07 Gb) and the least contiguous among the three both at the scaffold and contig level (scaffold/contig N50: 5.2Mb/30kb, NCBI accession number: GCA_000277835.1).

In all three clades, it has been suggested that shared genetic variation between (sub)-species within clades resulted from incomplete lineage sorting of ancestral polymorphisms, regardless of whether populations were connected by recent gene flow or not (Lamichhaney et al. 2015; Burri et al. 2015; Vijay et al. 2016). However, shared polymorphism is highly unlikely among clades because of their phylogenetic distance. Phylogenetic relationships and divergence time estimates between representatives of all three clades and zebra finch (*Taenopygia guttata*) as shown in **Figure 1** have been extracted as consensus of 10,000 phylogenetic reconstructions from Jetz et al. (2012, 2014) using the tree of 6670 taxa with sequence information by Ericson et al. (2006) as backbone (http://birdtree.org/). This places the separation between Corvoidea (crows) and Passerida (Darwin's finches, flycatcher) at over 50 million years corresponding to at least 8-25 million generations assuming a range in generation time between six years for hooded crows (Vijay et al. 2016), five years for Darwin's finches (Grant & Grant 1992) and two years for flycatchers (Brommer et al. 2004). With an estimated long-term N_e_ of 200,000 for flycatchers and crows (Wolf et al. 2010; Nadachowska-Brzyska et al. 2013; Vijay et al. 2016) and considerably less for Darwin's finches (Ne= 6,000 to 60,000, (Lamichhaney et al. 2015)) this yields a minimum range of 40-125 N_e_ generations as time to the most common ancestor. Since this is clearly beyond the expected time for complete lineage sorting (9-12 N_e_ generations; (Hudson et al. 2002)), species among the two clades are thus not expected to share ancestral polymorphism. The same consideration holds for the split between flycatcher and Darwin's finches assuming approximately 45 million years of divergence (**Fig. 1**). Even assuming an earlier, minimal age estimate of the split between Corvoidea and Passerida in the order of 25 million years ago (Jarvis et al. 2014; Prum et al. 2015; Jønsson et al. 2016) and a split between flycatchers and finches at 19 million years (Singhal et al. 2015) suggests complete lineage sorting for neutral variation with split times beyond 12 N_e_ generations.

### Establishing homology among genomes

Homologous regions between genomes were identified in order to quantify the degree to which genetic diversity, recombination, differentiation and divergence landscapes are conserved between species. To ensure comparability across all three clades in the most efficient way, we chose to lift-over coordinates of 50 kb non-overlapping windows from the genomes to the independent, well maintained high quality zebra finch reference genome (Hubbard 2002) that is closely related to all three clades. This approach assumes a high degree of synteny among species, which is justified given the evolutionary stasis of chromosomal organisation in birds across more than 100 million years of evolution (Ellegren 2010). Performing a base by base lift over can lead to partial loss of regions within a window as well as merging of non-adjacent windows. To avoid such errors we estimated the statistics for each species in windows prior to the lift over. Converting the coordinates of genomes from multiple different species into one single coordinate system allows for straightforward comparison of all statistics derived from the original polymorphism data (vcf files).

Whole genome alignments between species can be represented in the form of chain files that record the links between orthologous regions of the genome. We downloaded chain files from the UCSC website (https://genome.ucsc.edu/) to transfer the coordinates in bed format from flycatcher and Darwin's Finch genomes onto the zebra finch genome using the program liftOver (Kuhn et al. 2007). For the crow genome where no chain files were available, we first aligned the crow genome to the flycatcher genome using LASTZ (Harris 2007) to obtain a .psl file which was subsequently converted to a chain file using JCVI utility libraries (Tang et al. 2015). This chain file was then used to transfer the crow coordinates to zebra finch coordinates (via flycatcher) using the liftOver utility (Hinrichs et al. 2006).

Orthology could be established for a large proportion of the original genomes. Depending on parameter settings controlling stringency ('minmatch') and cohesion ('minblocks') percent recovery ranged from as little as 13% to over 90% (**Fig. S1, Table S1**). To find an optimal combination of parameter values and to validate liftOver quality, we made use of the fact that GC content in ortholgous regions of avian genomes are expected to be strongly conserved across long evolutionary distances (Weber et al. 2014). We calculated GC content in 50Kb windows from the three different assemblies and compared these values to the GC content at the new, orthologous positions lifted over to the zebra finch genome. Pearson's correlations were high across a broad set of parameter values in all clades ranging from 0.83-0.97. While the liftOver step is able to transfer the coordinates from the focal genome onto positions along the zebra finch genome these new positions do not retain the window structure from the original genomes. To be able to compare population genetic summary statistics between species in orthologous windows, we defined 50Kb windows along the zebra finch genome. For each window we then calculated a mean value across all lifted over regions that overlapped a given window. To ensure that this procedure of calculating means did not unduly influence comparability across species, we compared the values of GC content from each of the focal genomes after taking the mean across overlapping regions to the GC content in the zebra finch genomic windows. Although correlation coefficients were lower than those seen directly after the liftOver, they still exceeded 0.78, 0.82, 0.82 for Darwin's finch, flycatcher and crow respectively across a broad 'minmatch' and 'minblock' parameter space (**Fig. S1, Table S1**). The high correlation of GC content across the liftOver steps suggests that the lift over procedure of moving the windows from one genome assembly to another was reliable at the window size being evaluated. Finally, an optimal combination of stringency, cohesion and percent recovery was chosen on the basis of the (visually inferred) inflection point of the relationship between GC correlation and recovery (**Fig. S1)**.

It could be seen that certain regions of the genome were systematically more susceptible to drop out during liftOver than others for all clades (**Fig. S2**). I particular, regions located on scaffolds that have not been linked to any specific chromosome and those that have not been placed at a particular position along a chromosome were harder to liftOver than other regions of the genome. Hence, for the purpose of this study we have excluded these regions in all subsequent analyses. To ensure that the liftOver step did not introduce a large bias in the regions being analysed, we compared the GC content distribution of the regions that could be lifted over at different values of the "minmatch" parameter (**Fig. S3**). No clear evidence of bias with regard to GC content of the successfully lifted over regions emerged.

**Figure 2.**
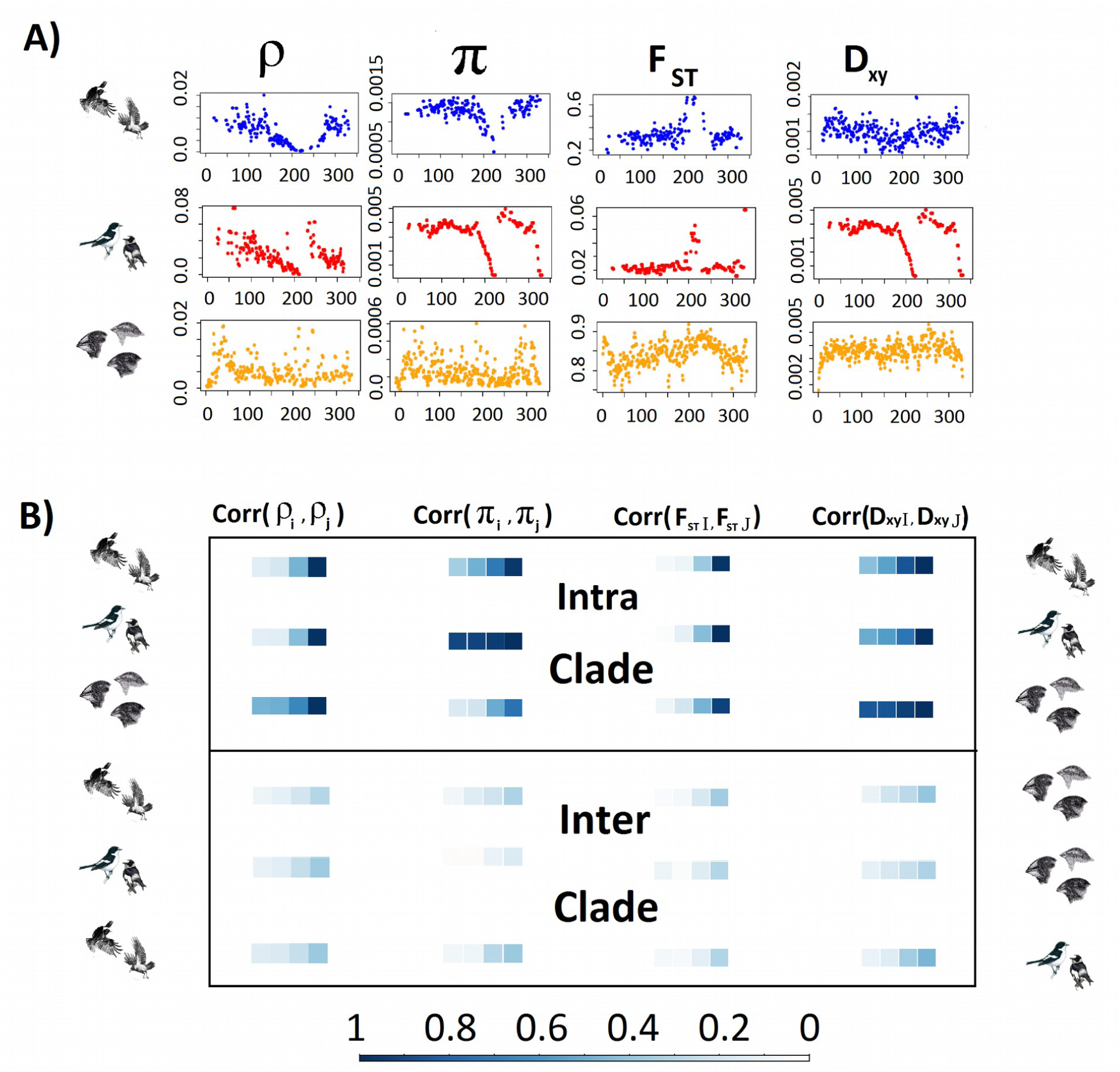
**Co-variation of population genetic summary statistics within and among clades A)** Genome-wide landscapes of four summary statistics are compared within and between clades. Depicted is an example showing the population recombination rate (ρ), nucleotide diversity (π), genetic differentiation (F_ST_) and sequence divergence (D_xy_) along chromosome 13 of zebra finch. The x-axis is scaled in units of 50kb windows. **B)** Distribution of correlation coefficients (Pearson's r) tabulated in 4 bins for population summary statistics characterizing variation within (ρ, π) and between populations (F_ST_, D_xy_). Correlations are first shown for population comparisons within each of the three clades (intra-clade). Subscripts *i, j* symbolize all possible combinations of correlations between two populations *i=1…n* and *j=i+1….n* for within-populations measures; Capital letters *I, J* symbolize inter-population statistics. Correlations exclude pseudo-replicated population comparisons. Similarly, within and between population measures were compared among all three clades, as illustrated by the bird images. In case of no association a Normal distribution centred around null would be expected.

**Figure 3.**
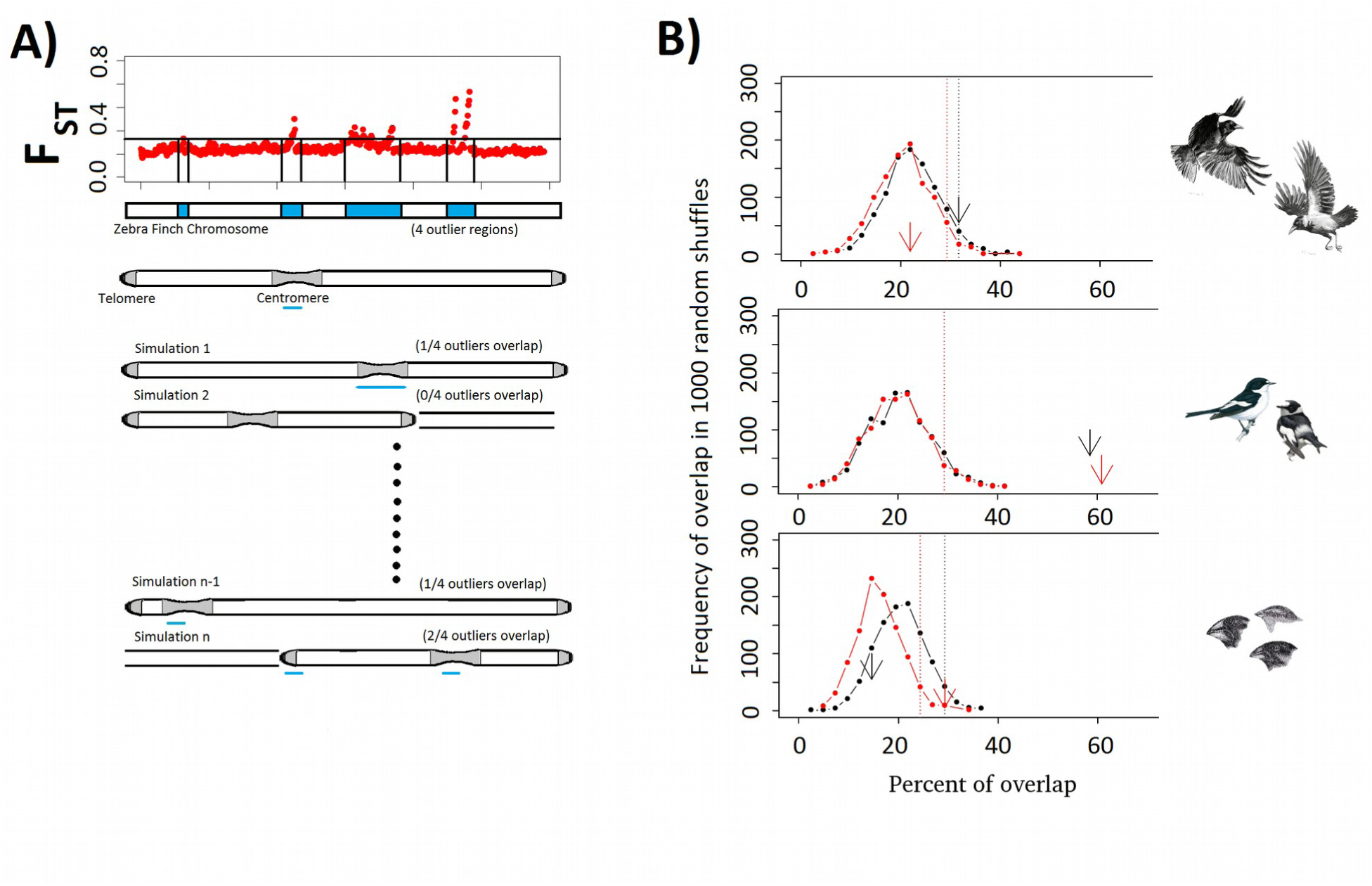
**Association of genomic differentiation landscapes with chromosomal features A)** Schematic of the shuffling of centromere and telomere positions to estimate the random expectation of the overlap. **B)** The degree of overlap between regions of elevated differentiation with regions adjacent to centro- and telomeres is quantified for two selected population pairs (red and black arrows) from each taxon and contrasted to distributions of random expectation as assessed by permutation.

### Datasets

We compiled the following population re-sequencing datasets for the three clades presented by order of divergence from zebra finch (**Table S2**). Populations with less than three individuals were excluded in all species.

1. Crows in the genus *Corvus* (124 genomes resequenced, 55 population comparisons within and between 2 focal species, the American crow *C. brachyrhynchos* and various (sub)-species and populations within the *C. (c.) spp.* complex). Population genetic summary statistics including genetic diversity (π), population recombination rate (ρ), genetic differentiation (F_ST_) and sequence divergence (D_xy_) across the European crow hybrid zone have been characterised using high coverage whole genome re-sequencing data of 60 individuals samples in a 2×2 population design between carrion crows *(Corvus (corone) corone)* and hooded crows *(C. (c.) cornix)* (Poelstra et al. 2014). This study has been followed by a broader sampling regime with a total of 118 crows from the *Corvus (c.) spp.* species complex including a parallel hybrid zone in Russia between *C. (c.) cornix* and *C. (c.) orientalis*, a contact zone between the latter and *C. (c.) pectoralis* and numerous other allopatric populations (Vijay et al. 2016). The system is still relative young. 12% of segregating genetic variation is still shared between Eurasian and American crows (*C. brachyrhynchos)* (Vijay et al. 2016) which split at approximately 3 million years ago (Jønsson et al. 2016). F_ST_ and D_xy_ ranged from 0.016-0.486 and 0.0015-0.0018 respectively. A broad range in π (0.0010-0.0033) and Tajima's D (0.5895 to -1.974) suggests perturbation by population specific demographic histories.
2. *Ficedula flycatchers* (200 genomes resequenced with 30 population comparisons across the 4 focal species *F. albicollis, F. hypoleuca, F. semitorquata and F. speculigera).* Species diverged approximately 2 million years ago and populations differ slightly in genome-wide levels of differentiation (π: 0.0029-0.0039). A total of 30 population comparisons within and across species provide a broad contrast across a spectrum of genome-wide differentiation (F_ST_: 0.012-0.981) and divergence (D_xy_: 0.0031-0.0050)(see (Burri et al. 2015)).
3. *Darwin's finches* (120 genomes resequenced, 44 population comparisons across the 6 focal species *G. conirostris, G. difficilis, C. pallidus, C. fusca, C. olivacea* and *P. inornata).* The differentiation landscape of Darwin's finches has been studied using whole genome re-sequencing data and has been instrumental in the identification of adaptive loci associated with beak shape evolution (Lamichhaney et al. 2015). This set of populations across several species differs slightly in genome-wide levels of diversity (π: 0.0003-0.0012, see (Lamichhaney et al. 2015). Species are estimated to share common ancestry ∼1.5 million years ago, yielding 44 population comparisons ranging across a broad spectrum of genome-wide differentiation (F_ST_: 0.192-0.897) and divergence (D_xy_: 0.0022-0.0047).

### Genetic diversity data

In all three study systems segregating genetic variation and related summary statistics have been characterized in non-overlapping windows across the genome using similar strategies based on the Genome Analysis Toolkit GATK (DePristo et al. 2011)(see individual studies for details). We used the final set of variant calls from each individual to calculate a set of summary statistics. vcf files were obtained from Lamichaney et al. (2015) for Darwin's finches, (2015) for flycatchers and (2016) for crows. Each of the statistics were calculated in 50 kb windows for all scaffolds longer than 50kb.

#### Population recombination rate (ρ) and nucleotide diversity (π)

To generate an estimate of the population scaled recombination rate in Darwin's finches ρ we followed the approach described in (Vijay et al. 2016). In brief, we used LDhelmet (Chan et al. 2012) on genotype data phased with fastPHASE (Scheet & Stephens 2006). The required mutation matrix was approximated from zebra finch substitution rates following Singhal et al. (2015). Population recombination rate data for crows and flycatchers were estimated using the same approach and were extracted from (Vijay et al. 2016) and (Kawakami et al. submitted), respectively. Pairwise nucleotide diversity π was calculated from the VCF files using the R package Hierfstat.

#### Genetic differentiation F_ST_, PBS and net divergence (D_XY_)

*F_st_* was estimated using Weir and Cockerham's estimator based on genotypes from the VCF files using the procedure implemented in the HIERFSTAT package as the ratio of the average of variance components. To avoid pseudo-replicated populations comparisons we also calculated lineage specific F_ST_ in the form of population branch statistics (PBS) using the formula PBS_Pop1=(-log(1-F_ST_(Pop1_Pop2)+(-log(1-F_ST_(Pop1_Pop3)))-log(1-F_ST_(Pop2_Pop3)/2. Custom scripts (Poelstra & Vijay 2014) that used the R package Hierfstat (Goudet 2005) were used to estimate *net divergence D_XY_.*

#### Quantifying similarity of genomic landscapes within and among clades

We used Pearson correlations as a simple means to characterize the degree of co-variation genome-wide distribution patterns for a given summary statistic. Correlation coeffecients were calculated on the basis of homologous windows within and between clades (see above). For intra-population measures (ρ, π) we calculated all possible combinations between two populations *i=1…n* and *j=i+1…n;* for inter-population metrics (F_ST_, PBS, D_XY_) all possible combinations between population comparisons *I, J.* Correlations exclude pseudo-replicated population comparisons (e.g. I=popA vs. popB, J=popA vs. popC). This yields a distribution of correlations coefficients for each summary statistic. Significance in co-variation between populations or population comparisons attributed if more than 95% of the distribution were above zero (significant positive correlation) or below zero (significant negative correlation).

#### Overlap with centromeres and telomeres

LiftOvers to the zebra finch genome in principle allow associating outlier regions from genome scans (e.g. islands of elevated differentiation) with genomic features such as centromeres or telomeres. This approach works under the assumption of karyotype conservation across large evolutionary timescales (Ellegren 2010). It is conservative in that overlap is only expected if centromere position is conserved between zebra finch and the taxon under consideration. Evolutionary lability of these features, partly expected due to known lineage-specific inversions in zebra finch (Romanov et al. 2014; Kawakami et al. 2014; Hooper & Price 2015) would reduce any real correlation (Type II error), but not introduce any spurious correlation (Type I error). Centromere and telomere positions were obtained for 22 and 20 chromosomes, respectively, in zebra finch from Knief & Forstmeier (2015). Regions identified as centromeres were on average ∼1Mb long (mean: 960,100 bp; range: 150,000 bp to 5,350,000 bp) while the sub-telomeric regions were shorter (mean: 169800; range: 50,000 bp to 298,700 bp). Some of the sub-telomeric regions and centromeres were located at the extreme ends of the chromosomes and orthologus regions could not be identified in the draft assemblies of the crow, flycatcher and Darwin's finch. These regions are either not assembled in the draft genomes or could not be lifted over unambiguously.

Of the 42 regions that have been identified as centromeres or sub-telomeric regions in zebra finch, orthologous regions could be identified for a subset of 38 in the flycatcher (mean recovery: 0.69), 39 in crow (mean recovery: 0.83) and only 25 in the Darwin's Finch genome (mean recovery: 0.55). The relatively low recovery in Darwin's finch is most likely owing to the lower quality of its genome, which is more highly fragmented than flycatcher and particularly crow. The telomeres of chromosome 5, 13 and 21 could not be lifted over in neither crow nor flycatcher genomes suggesting a systematic bias for these regions. To reduce the effect of such bias, we not only looked for overlap of outlier peaks (as defined below) with centromeres or telomeres, but also for overlap with increasing distance from the inferred positions of these features in five incremental steps of 10 kb. In case of random association no relationship would be expected with distance, in case of genuine association significance of the overlap should decrease with distance.

To relate characteristics of the genomic differentiation landscape to chromosomal features, we proceeded as follows. For each taxon we chose two independent population comparisons with the highest genome-wide average F_ST_ values. This strategy is owing to the fact that clear 'background peaks' caused by shared linked selection only start crystallising at an advanced level of population divergence (Burri et al. 2015; Vijay et al. 2016). This is theoretically expected and has been shown in crows where an increase in genome-wide F_ST_ is accompanied by an increase in autocorrelation between windows, peak overlap and the degree of co-variation in differentiation landscapes. Population pairs used and their corresponding differentiation statistics are shown in **Table S3**.

We then used positions along the zebra finch genome to calculate the percent of centromeres and telomeres that overlapped with differentiation outliers (**Table S4**). To check if the percent of overlap we observed was more than that expected by chance, we permuted the positions of centromeres and telomeres within each chromosome 1000 times using the shuffle option in bedtools (Quinlan & Hall 2010) and calculated the percent of overlap that was expected by chance alone. A significant association is inferred at type I error levels of 0.05/ 0.01 if the test statistic derived from the empirical centromere/telomere distribution exceeded a maximum of 4/0-times by test statistics derived from the permuted distributions.

## RESULTS

### Co-variation among populations within clades (micro-level)

It has been demonstrated for both flycatchers (Kawakami et al. submitted; Burri et al. 2015) and crows (Vijay et al. 2016) that summary statistics of genetic variation within and between populations were significantly correlated among populations. Extending the estimation of ρ, π, F_ST_, PBS and D_xy_ to the Darwin's finch complex corroborates the generality of this finding. Genome-wide patterns of all summary statistics were positively correlated among all populations in each of the three clades (**Fig. 2B, Table S5**). For ρ, correlation coefficients were highest in flycatchers (mean r=0.43), followed by Darwin's finches (*r* = 0.27) and crows (*r* = 0.19). Nucleotide diversity π showed strongest co-variation in flycatchers (r=0.95), followed by crows (r=0.70) and Darwin's Finches (r=0.49). *F_ST_* correlations were consistently positive between all population pairs in Darwin's finches (r=0.46), flycatchers (mean r=0.42) and crows (r=0.36). Correlation of genetic differentiation patterns was even stronger when considering the lineage-specific population branch statistic (PBS). *D_XY_* similarly showed exclusively positive correlations within clades with mean correlation coefficients of 0.72, 0.85 and 0.94 between flycatcher, crow and Darwin's finch populations, respectively. Importantly, it was negatively correlated with F_ST_(mean range r=-0.19 to -0.45). This is predicted by long-term linked selection (acting already in the ancestor) and opposed to the expectation for divergent selection against gene flow (Nachman & Payseur 2012; Cruickshank & Hahn 2014). Overall, these results support a contribution of linked selection unrelated to population-specific selection, as has been previously suggested for crows (Vijay et al. 2016) and using independent recombination data also for flycatchers (Kawakami et al. submitted; Burri et al. 2015).

### Co-variation among populations across clades (macro-level)

Next, we investigated whether summary statistics indicative of local N_e_ also co-varied in syntenic regions across clades precluding any influence of shared history in demographic or selective processes (**Supporting Information**). Though effect sizes were lower, correlations were consistently positive for all summary statistics. This was true for comparisons between flycatcher and Darwin's finch populations, as well as for the more divergent comparisons between flycatchers/Darwin's finches and crow, representing 50 million years of evolution (**Fig. 2, Table 1, Fig. S4).** As in the micro-level comparisons, D_XY_ and F_ST_ were negatively correlated among clades (mean range r=-0.21 to -0.16).

**Table 1:** Correlations between clades for the population-scaled recombination rate (ρ), nucleotide diversity (л), genetic differentiation (F_ST_, PBS), sequence divergence (D_xy_). *Abbreviations*: CR = crow, FC = flycatcher, DF=Darwin's finch.

| Clade | Statistics | Minimum | Lower 5% | Mean | Max |
| --- | --- | --- | --- | --- | --- |
| CR vs DF | $\rho$ | 0.010 | 0.025 | 0.102 | 0.239 |
| FC vs DF |  | 0.073 | 0.089 | 0.172 | 0.256 |
| CR vs FC |  | 0.008 | 0.028 | 0.099 | 0.192 |
| CR vs DF | $\pi$ | 0.056 | 0.132 | 0.204 | 0.301 |
| FC vs DF |  | -0.015 | -0.011 | 0.082 | 0.149 |
| CR vs FC |  | 0.057 | 0.060 | 0.271 | 0.364 |
| CR vs DF | $F_{ST}$ | 0.020 | 0.041 | 0.163 | 0.341 |
| FC vs DF |  | 0.078 | 0.014 | 0.138 | 0.287 |
| CR vs FC |  | 0.044 | 0.038 | 0.115 | 0.293 |
| CR vs DF | PBS | 0.075 | 0.108 | 0.185 | 0.283 |
| FC vs DF |  | 0.091 | 0.122 | 0.231 | 0.353 |
| CR vs FC |  | 0.136 | 0.150 | 0.223 | 0.361 |
| CR vs DF | $D_{xy}$ | 0.087 | 0.197 | 0.268 | 0.387 |
| FC vs DF |  | 0.100 | 0.160 | 0.224 | 0.306 |
| CR vs FC |  | 0.089 | 0.131 | 0.342 | 0.454 |

### Overlap with structural genomic features

We next sought to investigate the potential impact of structural genomic features where the effect of linked selection might be particularly pronounced. We evaluated whether regions of highly elevated differentiation were associated with regions of suppressed recombination adjacent to centromeres and subtelomeric regions as predicted from the location of such regions in zebra finch (karyotype data is lacking for both crow and collared flycatcher; **Fig. 3A**). For each clade we focused on the two most divergent population/species comparisons (Burri et al. 2015; Vijay et al. 2016). In all three clades, overlap was significantly greater than expected by chance in at least one comparison of each species (percentage of overlap in flycatchers: 58.53% and 60.98%, crows: 21.95% and 31.7%, Darwin's finches: 14.63% and 29.27%) (**Fig. 3B**). Considering regions next to centromeres and sub-telomeric regions separately suggested significant association for subtelomeric regions in all three clades (**Fig. S5**), for centromeres only in flycatcher (**Fig. S6**).

## DISCUSSION

In a comparative approach we leveraged information from genome-wide patterns of genetic diversity and differentiation shared across micro- and macro-evolutionary timescales. We used multiple population samples of three distantly related avian clades with split times beyond the expected time for complete lineage sorting. Genome-wide heterogeneity in genetic variation captured by population genetic statistics reflective of regional N_e_ co-varied among clades across 50 million years of evolution. This finding supports a role of linked selection unrelated to selection against gene flow, the latter promoting adaptation and/or reproductive isolation in a population specific context. The degree of correlation within, but in particular among clades is remarkable considering divergence times of several million generations, gaps in syntenic alignments and the statistical error generally associated with population genetic estimates. Interestingly, the magnitude of correlations was not related to divergence time (**Fig. S4**) with sometimes noticeably higher correlation coefficients for the phylogenetically older flycatcher-crow comparison, than for the younger flycatcher-finch comparison (**Table 1**). This suggests that the strength of co-variation may be underestimated by factors such as genome quality (fragmented in Darwin's finch), population sampling (lower sample size for Darwin's finch) and/or differences in the degree of rearrangements between clades

In addition to linked selection, evolutionary stable variation in regional mutation rate may impact diversity levels at equilibrium (θ=4N_e_µ) among clades in a similar fashion. However, genetic diversity is generally only weakly associated with mutation rate (Cutter & Payseur 2013; Vijay et al. 2016) Instead, genetic diversity shows a strong positive relationship with recombination rate (Nachman & Payseur 2012; Burri et al. 2015). With little evidence for recombination-associated mutation (and hence r∼µ) (Cutter & Payseur 2013) linked selection reducing regional genomic levels of N_e_ as a function of recombination rate appears to be an important candidate process underlying genomic variation in levels of genetic diversity (Cutter & Payseur 2013; Cruickshank & Hahn 2014; Slotte 2014). Our results demonstrating co-varying diversity and differentiation landscapes across clades are suggestive of persistence in linked selection at syntenic genomic regions. Moreover, the observation that sequence divergence (D_xy_) was generally reduced in areas of high relative differentiation (F_ST_, PBS) both within and across clades further points towards a selective process continuously purging diversity and reducing effective population size already in ancestral populations (Cruickshank & Hahn 2014).

Linked selection can occur in the form of background selection (Charlesworth 1994) or recurrent hitch-hiking dynamics by selective sweeps (Smith & Haigh 1974). Both processes reduce genome-wide diversity and leave signatures of selection that are difficult to discern (Stephan 2010). Consistent with both, recent population genetic studies of flycatchers and crows suggest that diversity and differentiation landscapes were associated with variation in recombination rate and gene density (as a proxy for target for selection) (Burri et al. 2015; Vijay et al. 2016). In a model based approach Corbett-Detig (2015) assessed the relative likelihood of both processes concluding that for species with low/moderate population sizes (including flycatchers) background selection would prevail over hitch-hiking in relative importance. Here, we also found regions of reduced diversity and elevated differentiation to be associated with candidate centromeric regions as lifted over from zebra finch. This overlap, suggested also for other systems (Delmore et al. 2015; Roesti et al. 2015) may be a consequence of lower recombination rates at centromeres promoting background selection. It is puzzling, however, that we even found overlap at regions adjacent to telomeres that are not necessarily characterized by low recombination in birds (Backström et al. 2010; Kawakami et al. 2014). This tentatively suggests that in addition to background selection recurrent positive selection might be at play. Further evaluation of this hypothesis will require fine-scale recombination rate estimates across all clades, better assemblies of centromeric and sub-telomeric regions, a map of their exact location, and on the bioinformatic side improved methods for translating genomic coordinates among distantly related species.

Using a comparative approach we shed light on the processes mediating a similar landscape of heterogeneity in diversity and differentiation across large evolutionary timescales. We advocate increased use of comparative, phylogenetic approaches to understand the processes underlying signatures of selection to help interpret results from commonly conducted genome scans.

## Data Accessibility

Raw data forming the basis for this study are publicly available at PRJNA192205 & PRJEB9057 (Crows), PRJEB2984 (Flycatchers), PRJNA301892 (Darwin's Finches).

## Author Contributions

NV and JW conceived of the study, NV conducted all bioinformatic analyses with help from MW. RB, TK and HE provided population genetic summary statistics for the flycatcher. NV and JW wrote the manuscript with input from all other authors.

## ACKNOWLEDGMENTS

Funding for this study was provided by the Swedish Research Council (grant number 621-2010-5553 to J.W. and 2014-6325 to T.K. and 2013-08721 to H.E.), Marie Sklodowska Curie Actions (grant number 600398 to T.K.), the European Research Council (grant number ERCStG-336536 to J.W.), the Knut and Alice Wallenberg Foundation (to H.E.) and the Swiss National Science Foundation (grants number PBLAP3-134299 and PBLAP3_140171 to R.B.). We are grateful for the access to the computational infrastructure provided by the UPPMAX Next-Generation Sequencing Cluster and Storage (UPPNEX) project, funded by the Knut and Alice Wallenberg Foundation and the Swedish National Infrastructure for Computing. We would like to thank Leif Andersson and his group for providing access to the genotype data from Lamichhaney et al. (2015).

## REFERENCES

1 Auton A,Fledel-Alon A, Pfeifer S et al. (2012) A fine-scale chimpanzee genetic map from population sequencing. Science, 336, 193–198.

2 Backström N, Forstmeier W, Schielzeth H et al. (2010) The recombination landscape of the zebra finch Taeniopygia guttata genome. Genome Research, 20, 485–495.

3 Bank C, Ewing GB, Ferrer-Admettla A, Foll M, Jensen JD (2014) Thinking too positive? Revisiting current methods of population genetic selection inference. Trends in Genetics, 30, 540–546.

4 Brommer JE, Gustafsson L, Pietiäinen H, Merilä J (2004) Single‐generation estimates of individual fitness as proxies for long‐term genetic contribution. The American Naturalist, 163, 505–517.

5 Burri R, Nater A, Kawakami T et al. (2015) Linked selection and recombination rate variation drive the evolution of the genomic landscape of differentiation across the speciation continuum of Ficedula flycatchers. Genome Research, 25:1656–1665.

6 Chan AH, Jenkins PA, Song YS (2012) Genome-wide fine-scale recombination rate variation in Drosophila melanogaster PLoS Genetics, 8, e1003090.

7 Charlesworth B (1994) The effect of background selection against deleterious mutations on weakly selected, linked variants. Genetical Research, 63, 213–227.

8 Charlesworth B (1998) Measures of divergence between populations and the effect of forces that reduce variability. Molecular Biology and Evolution, 15, 538–543.

9 Charlesworth B, Morgan MT, Charlesworth D (1993) The effect of deleterious mutations on neutral molecular variation. Genetics, 134, 1289–1303.

10 Corbett-Detig RB, Hartl DL, Sackton TB (2015) Natural selection constrains neutral diversity across awide range of species. PLoS Biol, 13, e1002112.

11 Cruickshank TE, Hahn MW (2014) Reanalysis suggests that genomic islands of speciation are due to reduced diversity, not reduced gene flow. Molecular Ecology, 23, 3133–3157.

12 Cutter AD, Payseur BA (2013) Genomic signatures of selection at linked sites: unifying the disparity among species. Nature Reviews. Genetics, 14, 262–274.

13 Delmore KE, Hübner S, Kane NC et al. (2015) Genomic analysis of a migratory divide reveals candidate genes for migration and implicates selective sweeps in generating islands of differentiation. Molecular Ecology, 24, 1873–1888.

14 DePristo MA, Banks E, Poplin R et al. (2011) A framework for variation discovery and genotyping using next-generation DNA sequencing data. Nature Genetics, 43, 491–498.

15 Dettman JR, Sirjusingh C, Kohn LM, Anderson JB (2007) Incipient speciation by divergent adaptation and antagonistic epistasis in yeast. Nature, 447, 585–588.

16 Ellegren H (2010) Evolutionary stasis: the stable chromosomes of birds. Trends in Ecology & Evolution, 25, 283–291.

17 Ellegren H, Galtier N (2016) Determinants of genetic diversity. Nature Reviews Genetics, 17, 422–433.

18 Ellegren H, Smeds L, Burri R et al. (2012) The genomic landscape of species divergence in Ficedula flycatchers. Nature, 491, 756–760.

19 Ericson PGP, Zuccon D, Ohlson JI et al. (2006) Higher-level phylogeny and morphological evolution of tyrant flycatchers, cotingas, manakins, and their allies (Aves: Tyrannida). Molecular Phylogenetics and Evolution, 40, 471–483.

20 Goudet J (2005) hierfstat, a package for R to compute and test hierarchical F-statistics. Molecular Ecology Notes, 5, 184–186.

21 Grant PR, Grant BR (1992) Demography and the genetically effective sizes of two populations of Darwin’s finches. Ecology, 73, 766–784.

22 Harris RS (2007) Improved pairwise alignment of genomic DNA. PhD thesis Pennsylvania State University

23 Hinrichs AS, Karolchik D, Baertsch R et al. (2006) The UCSC genome browser database: update 2006. Nucleic Acids Research, 34, D590–D598.

24 Hodgkinson A, Eyre-Walker A (2011) Variation in the mutation rate across mammalian genomes. Nature Reviews Genetics, 12, 756–766.

25 Hooper DM, Price TD (2015) Rates of karyotypic evolution in Estrildid finches differ between island and continental clades: chromosome inversion in finches. Evolution, 69, 890–903.

26 Hubbard T (2002) The Ensembl genome database project. Nucleic Acids Research, 30, 38–41.

27 Hudson RR, Coyne JA, Huelsenbeck J (2002) Mathematical consequences of the genealogical species concept. Evolution, 56, 1557–1565.

28 Jarvis ED, Mirarab S, Aberer AJ et al. (2014) Whole-genome analyses resolve early branches in the tree of life of modern birds. Science, 346, 1320–1331.

29 Jetz W, Thomas GH, Joy JB et al. (2014) Global distribution and conservation of evolutionary distinctness in birds. Current Biology, 24, 919–930.

30 Jetz W, Thomas GH, Joy JB, Hartmann K, Mooers AO (2012) The global diversity of birds in space and time. Nature, 491,444–448.

31 Jønsson KA, Fabre P-H, Kennedy JD et al. (2016) A supermatrix phylogeny of corvoid passerine birds (Aves: Corvides). Molecular Phylogenetics and Evolution, 94, Part A, 87–94.

32 Kawakami T, Mugal CF, Suh A et al. (submitted) Whole-genome patterns of linkage disequilibrium in flycatcher genomes clarify the causes and consequences of fine-scale recombination rate variation in birds.

33 Kawakami T, Smeds L, Backström N et al. (2014) A high-density linkage map enables a second-generation collared flycatcher genome assembly and reveals the patterns of avian recombination rate variation and chromosomal evolution. Molecular Ecology, 23, 4035–4058.

34 Knief U& WF (2015) Mapping centromeres of microchromosomes in the zebra finch (Taeniopygia guttata) using half-tetrad analysis. Chromosoma, 125:757–68

35 Kronforst MR, Papa R (2015) The functional basis of wing patterning in heliconius butterflies: the molecules behind mimicry. Genetics, 200, 1–19.

36 Kuhn RM, Karolchik D, Zweig AS et al. (2007) The UCSC genome browser database: update 2007. Nucleic Acids Research, 35, D668–73.

37 Lamichhaney S, Berglund J, Almén MS et al. (2015) Evolution of Darwin’s finches and their beaks revealed by genome sequencing. Nature, 518, 371–375.

38 Nachman MW, Payseur BA (2012) Recombination rate variation and speciation: theoretical predictions and empirical results from rabbits and mice. Philosophical Transactions of the Royal Society of London. Series B, Biological Sciences, 367, 409–421.

39 Nadachowska-Brzyska K, Burri R, Olason PI et al. (2013) Demographic divergence history of pied flycatcher and collared flycatcher inferred from whole-genome re-sequencing data. PLoS Genetics, 9, e1003942.

40 Nosil P, Feder JL (2013) Genome evolution and speciation: toward quantitative descriptions of pattern and process. Evolution, 67, 2461–2467.

41 Poelstra JW, Vijay N (2014) The genomic landscape underlying phenotypic integrity in the face of gene flow in crows. Science, 344, 1410–1414.

42 Poelstra JW, Vijay N, Hoeppner MP, Wolf JBW (2015) Transcriptomics of colour patterning and coloration shifts in crows. Molecular Ecology, 24, 4617–4628.

43 Prum RO, Berv JS, Dornburg A et al. (2015) A comprehensive phylogeny of birds (Aves) using targeted next-generation DNA sequencing. Nature, 526, 569–573.

44 Quinlan AR, Hall IM (2010) BEDTools: a flexible suite of utilities for comparing genomic features. Bioinformatics, 26, 841–2.

45 Rockman MV, Skrovanek SS, Kruglyak L (2010) Selection at linked sites shapes heritable phenotypic variation in C. elegans. Science, 330, 372–376.

46 Roesti M, Kueng B, Moser D, Berner D (2015) The genomics of ecological vicariance in threespine stickleback fish. Nature Communications, 6, 8767.

47 Romanov MN, Farré M, Lithgow PE et al. (2014) Reconstruction of gross avian genome structure, organization and evolution suggests that the chicken lineage most closely resembles the dinosaur avian ancestor. BMC Genomics, 15, 1060.

48 Scheet P, Stephens M (2006) A fast and flexible statistical model for large-scale population genotype data: applications to inferring missing genotypes and haplotypic phase. The American Journal of Human Genetics, 78, 629–644.

49 Seehausen O, Butlin RK, Keller I et al. (2014) Genomics and the origin of species. Nature Reviews Genetics, 15, 176–192.

50 Singhal S, Leffler EM, Sannareddy K et al. (2015) Stable recombination hotspots in birds. Science, 350, 928–932.

51 Slotte T (2014) The impact of linked selection on plant genomic variation. Briefings in Functional Genomics, 13, 268–275.

52 Smith JM, Haigh J (1974) The hitch-hiking effect of a favourable gene. Genetical Research, 23, 23–35.

53 Soria-Carrasco V, Gompert Z, Comeault AA et al. (2014) Stick insect genomes reveal natural selection’s role in parallel speciation. Science, 344, 738–742.

54 Stephan W (2010) Genetic hitchhiking versus background selection: the controversy and its implications. Philosophical Transactions of the Royal Society of London B: Biological Sciences, 365, 1245–1253.

55 Strasburg JL, Sherman NA, Wright KM et al. (2012) What can patterns of differentiation across plant genomes tell us about adaptation and speciation? Philosophical Transactions of the Royal Society of London B: Biological Sciences, 367, 364–373.

56 Tang H, Li J, Krishnakumar V (2015) jcvi: JCVI utility libraries.

57 Vijay N, Bossu CM, Poelstra JW, Weissensteiner MH, Suh A, Kryukov AP, Wolf JBW. (2016) Evolution of heterogeneous genome differentiation across multiple contact zones ina crow species complex. Nature Communications 7, 13195.

58 Weber CC, Boussau B, Romiguier J, Jarvis ED, Ellegren H (2014) Evidence for GC-biased gene conversion as a driver of between-lineage differences in avian base composition. Genome biology, 15, 549.

59 Wolf JBW, Bayer T, Haubold B et al. (2010) Nucleotide divergence versus gene expression differentiation: comparative transcriptome sequencing in natural isolates from the carrion crow and its hybrid zone with the hooded crow. Molecular Ecology, 19, 162–175.

60 Wolf JBW, Ellegren H (2016) Making sense of genome scans in the light of speciation. Nature Reviews Genetics, in press.

